# Prostaglandin E2 prevents radiotherapy-induced alopecia by attenuating transit amplifying cell apoptosis through promoting G1 arrest

**DOI:** 10.1101/2022.11.24.517788

**Authors:** Shih-Fan Lai, Wen-Yen Huang, Wei-Hung Wang, Jin-Bon Hong, Sung-Hsin Kuo, Sung-Jan Lin

## Abstract

Growing hair follicles (HFs) harbor actively dividing transit amplifying cells (TACs), rendering them highly sensitive to radiotherapy (RT). Clinically, there is still a lack of effective treatment for radiotherapy-induced alopecia (RIA). We aimed to dissect the effect and mechanism of local prostaglandin E2 (PGE2) pretreatment in RIA prevention. We found that PGE2 pretreatment reduced RIA by preventing premature termination of anagen through enhancing HF self-repair.

Mechanistically, PGE2 did not activate HF stem cells, but preserved more TACs for regenerative attempts. Pretreatment of PGE2 lessened radiosensitivity of TACs by transiently arresting them in the G1 phase, thereby reducing TAC apoptosis and mitigating HF dystrophy. The preservation of more TACs accelerated HF self-repair and bypassed RT-induced premature catagen entry. Promoting G1 arrest by systemic administration of palbociclib isethionate (PD0332991), a CDK4/6 inhibitor, offered a similar protective effect against RT. Therefore, PGE2 protects HF TACs from RT by transiently inducing G1 arrest, and the regeneration of HF structures lost from RT is accelerated to resume anagen growth, thus bypassing the long downtime of hair loss. PGE2 has the potential to be repurposed as a preventive treatment for RIA.

## Introduction

Radiotherapy (RT) with ionizing radiation (IR) is employed for the treatment of approximately 50% cancer patients [1]. Despite improvements in IR delivery technology, injury to normal tissues is still inevitable [2, 3] and radiotherapy-induced alopecia (RIA) is a common side effect[4, 5]. Hair follicles (HFs) undergo cyclic growth through the anagen, catagen, and telogen phases [6, 7]. Quiescent HF stem cells (HFSCs) located in the bulge epithelium are transiently activated in early anagen to fuel HF regeneration [7, 8]. In anagen, HF matrix cells, the transit amplifying cells (TACs) located in hair bulbs, actively divide to support continuous hair shaft elongation[6, 8, 9]. TACs of HFs are among the most proliferative cells in the body [10, 11], rendering anagen HFs highly susceptible to RT. Hair loss is almost unavoidable during RT for head and neck regions [4, 5, 12, 13]. Since hair, a key integral element of our appearance, is essential for social communication, RIA negatively impacts on cancer patients psychosocially [4, 5]. Methods to attenuate RIA are of high clinical significance.

RT causes HF dystrophy by inducing TAC apoptosis from IR-induced DNA breaks, resulting in premature catagen/telogen entry and hair loss [11, 14]. It takes approximately 3 months after RT for HFs to regrow from telogen back to anagen via activation of HFSCs [15]. When HFs are severely injured, hair regrowth is never adequate, and RIA becomes persistent or permanent [5, 12, 16]. Regrettably, there is still a lack of effective treatments to prevent RIA. We showed that anagen HFs are capable of eliciting a self-reparative attempt, named anagen HF repair (AHFR), to restore structures lost from IR injury by recruiting progenitor cells in or adjacent to the hair bulb to replenish TACs [11, 17]. In the face of more severe HF dystrophy from higher IR doses, it takes a longer time to activate HFSCs, and hair loss is more severe [11, 17]. Promoting AHFR by accelerating the recruitment of local progenitor cells to replenish TACs can attenuate hair loss from IR in animal models [11].

Prostanoids are a group of lipid mediators generated by cyclooxygenase metabolites of the arachidonic acid [18]. Prostanoids are widely generated in the body in response to diverse stimuli [18, 19]. Among them, prostaglandin E2 (PGE2) has been shown to be pivotal for tissue regeneration in multiple tissues [19, 20]. Previous results show the beneficial effects of PGE2 on tissue stem cells for tissue regeneration [21–24]. PGE2 given immediately after IR is able to reduce hematopoietic stem cell loss [21, 22]. In the intestine, PGE2 enhances intestinal crypt stem cell survival for the maintenance of epithelial homeostasis after IR injury [23, 25]. PGE2 was shown to be promising in protecting HFs from IR injury in animal models, but the mechanism is unclear [27, 28]. Clinically, PGE2 has long been used to facilitate labor progression [26]. Owing to its safety in clinical use, it can be repurposed for the prevention or treatment of RIA. However, how PGE2 works and how PGE2 should be administered to yield a better protective effect on HFs remain to be explored. Here, we investigated the effect and mechanism of local PGE2 pretreatment in RIA prevention.

## Materials and Methods

### Mice

All animal experiments were approved by the local institutional animal care and use committee. C57BL/6 mice were obtained from National Laboratory Animal Center, Taiwan. Fluorescent ubiquitination-based cell cycle indicator (FUCCI) mice [29] were obtained from the RIKEN BioResource Center (Kyoto, Japan). For invasive experiments, animals were anesthetized by intramuscular injection of tiletamine-zolazepam (Telazol^®^).

### Radiation exposure

The mouse dorsal hair was carefully shaved on postnatal day 32 (early full anagen). A single dose of γ-radiation (8.5Gy) was delivered from the dorsal side using a 137Cs source-662 keV photons (dose rate, 3.37 Gy/min; γ-irradiator IBL 637, CIS Bio International, Saclay, France).

### dmPGE2 treatment

dmPGE2 (Cayman Chemical, Ann Arbor, MI, USA) was prepared at 50 μg/mL. dmPGE2 (2 μg/g of body weight) divided into 4 doses for the 4 quadrants of the dorsal skin or vehicle was injected subcutaneously 2 hours before IR. Skin samples were harvested at the indicated time points for histological analysis. To label the proliferating cells, 5-bromo-20-deoxyuridine (BrdU) (50 mg/kg body weight) was injected intraperitoneally 1 hour before skin samples were harvested. Continuous BrdU labeling was performed to map the proliferative activity of HFSCs.

### Histology, immunostaining, and terminal deoxynucleotidyl transferase dUTP nick end labeling staining

Skin samples were fixed for paraffin embedding and sectioned (4–5μm thick) for hematoxylin and eosin, terminal deoxynucleotidyl transferase dUTP nick end labeling (TUNEL, Promega, Madison, WI, USA), immunohistochemistry (IHC), and immunofluorescence stainings. IHC and immunofluorescence stainings were performed with antigen retrieval, as suggested by the antibody manufacturers. The antibodies used are listed in **Supplementary Table S1**. Image acquisition, quantification, cell culture, time-lapse images, quantification of γ-H2AX foci and statistical analysis were summarized in the **Supplementary methods**.

## Results

### Local PGE2 pretreatment prevents IR-induced hair loss by converting a dystrophic catagen response to a dystrophic anagen response

In C57BL/6 mice, the first anagen starts at 4 weeks of age and persists for approximately 2 weeks before entering the catagen phase [30]. We used this window of synchronized anagen to investigate the effects of IR on anagen HFs. We previously showed that a single IR dose up to 5.5Gy could only induce a dystrophic anagen response in mouse dorsal HFs, whose structures are restored by an AHFR response [11, 17, 31]. To mimic RT-induced hair loss that is characterized by a dystrophic catagen response [15], we increased IR to 8.5Gy and found anagen HFs irradiated by 8.5Gy on postnatal day 32 prematurely initiated catagen and progressed to telogen on day 7 (**Fig.1A, 1B, Supplementary Fig.1**), consistent with a dystrophic catagen response [14, 15]. To test whether PGE2 could protect HFs from IR injury, we locally injected PGE2 2 hours before IR. The hair loss was markedly reduced (**Fig.1A**). Dorsal hair was retained on day 7 after IR. Histologically, the hair bulb shrank, and the HFs progressively shortened from 6 to 96 hours (**Fig.1B**). Then, the hair bulb enlarged again, and the HF progressively elongated from 96 hours to 7 days after IR. Although PGE2 pretreatment did not completely prevent HF dystrophy, it enabled HFs to successfully repair themselves. Therefore, PGE2 pretreatment prevented IR-induced hair loss by converting a dystrophic catagen response to a dystrophic anagen response.

**Figure 1.**
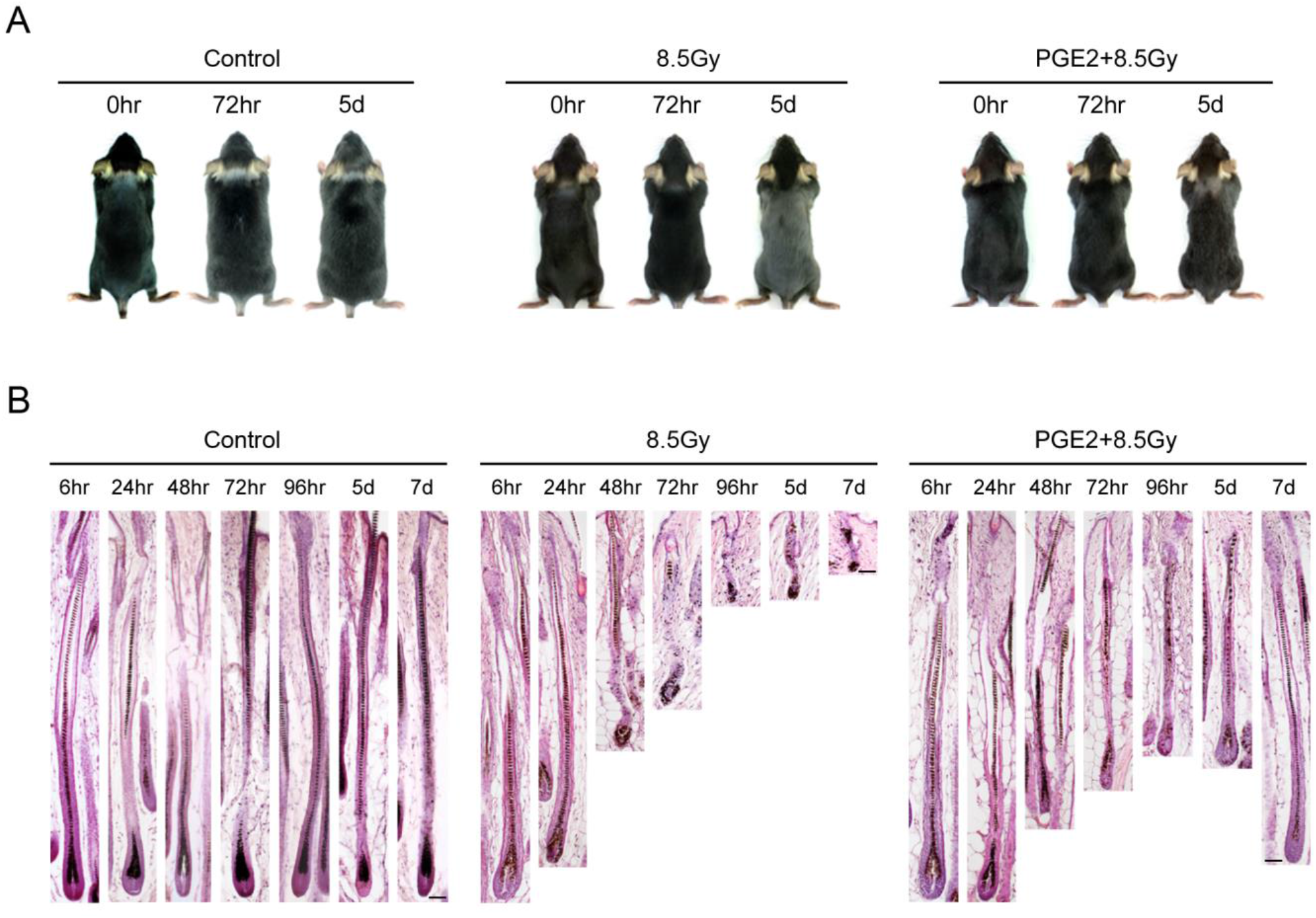
Local PGE2 pretreatment reduces hair loss from IR damage. **(A)** Gross pictures after IR. Prominent hair loss was induced by 8.5Gy of IR. With PGE2 pretreatment 2 hours before IR, hair loss was pronouncedly reduced. (**B**) Histology of serial changes of HFs. H&E staining. 8.5Gy of IR resulted in severely dystrophic changes of HFs with the induction of premature catagen entry. With local PGE2 pretreatment, HF dystrophy was lessened and the anagen HF structure was restored on day 7 post-IR. Scale bar =100 μ m in B.

### PGE2 pretreatment reduces cell death and increases compensatory cell proliferation after IR

To explore how PGE2 pretreatment protects HFs from IR-induced dystrophy, we first assessed cell proliferation using BrdU pulse labeling in PGE2-treated and untreated mice following IR. Without PGE2 pretreatment, cell proliferation in the hair bulb was halted 6 hours after IR (**Fig.2A**). There was a slight and transient restoration of cell proliferation in the hair bulb from 12 to 48 hours. Cell proliferation completely ceased at 72 hours, consistent with the progression of a dystrophic catagen response. In contrast, after PGE2 pretreatment, although cell proliferation in the hair bulb significantly decreased at 6 hours after IR (**Fig.2A**), cell proliferation increased again at 12 hours, with fluctuations from 24 to 72 hours. Cell proliferation steadily increased from 72 hours to 6 days after IR and was then maintained at a steady high level. This re-established steady cell proliferation correlated with a full anagen architecture re-established by a successful reparative attempt (**Fig. 1B**).

**Figure 2.**
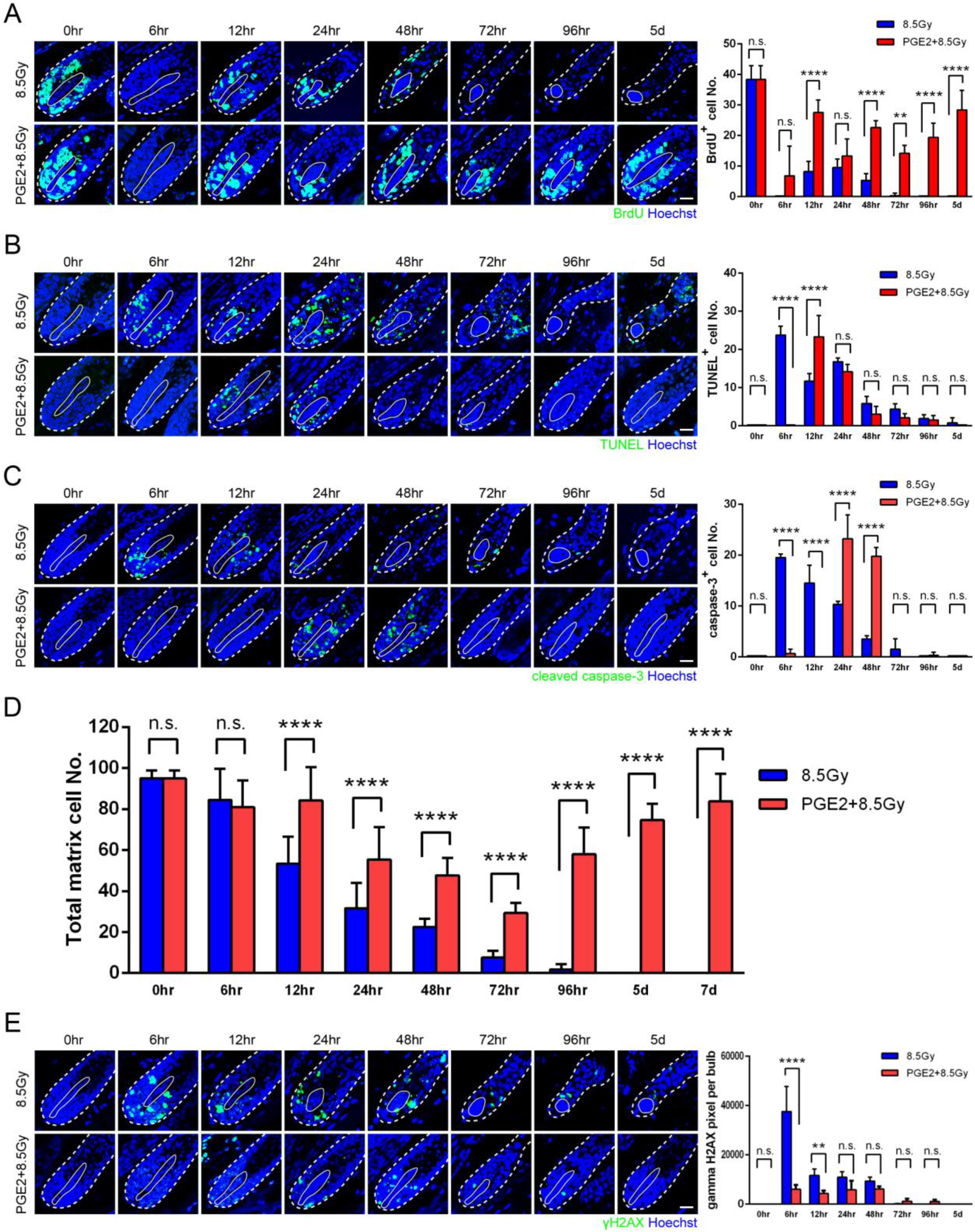
PGE2 pretreatment reduces IR-induced apoptosis of TACs. (**A**) Pulse BrdU labeling for analysis of cell proliferation in the hair bulb. BrdU positive cells in each hair bulb is quantified on the right side. (**B**) TUNEL staining for analysis of apoptosis. Quantification of TUNEL+ matrix cells in the hair bulb is shown on the right side. (**C**) Quantification of matrix cell numbers in the hair bulb. PGE2 pretreatment preserves a higher matrix cell number post-IR. (**D**) Immunofluorescent staining of cleaved caspase-3. Quantification of cleaved caspase-3+ matrix cells in the hair bulb is shown on the right side. (**E**) Immunofluorescent staining of γ-H2AX. Quantification of foci of γ-H2AX in each hair bulb is shown on the right side. Statistical significance was determined by Student’s *t* test. * *p* < 0.05, ** *p* < 0.01, *** *p* < 0.001, **** *p* < 0.0001. n.s.: non-significant. Error bars, mean + SEM. Dashed line: basement membrane. Solid line: dermal papilla. Scale bar = 20 μ m.

Next, we analyzed apoptosis in HFs using TUNEL and cleaved caspase-3 stainings [11]. Both assays revealed a similar trend **(Fig.2B,2C)**. Without PGE2 pretreatment, massive apoptosis occurred at 6 hours after IR, and then cell apoptosis decreased but persisted until day 5 (**Fig.2B**). In the PGE2 pretreatment group, apoptosis also occurred but was delayed. TUNEL staining was not positive until 12 hours after IR (**Fig.2B**), and TUNEL positive cells decreased from 24 to 96 hours. Compared with the non-PGE2-treated group, TUNEL positive matrix cells were significantly reduced from 6 to 12 hours (**Fig.2B**). Analysis of double-stranded DNA breaks showed γ-H2AX foci greatly increased at 6 hours after IR in the non-PGE2-treated group (**Fig.2E**). Following this peak, foci of γ-H2AX decreased from 12 to 96 hours. In contrast, γ-H2AX levels greatly decreased after PGE2 pretreatment. The foci of γ-H2AX were maintained at a low level from 6 to 96 hours after IR and then disappeared (**Fig.2E**).

We quantified and compared the number of hair matrix cells. Without PGE2 pretreatment, IR injury led to a progressive decrease of matrix cell numbers from 6 to 96 hours (**Fig.2D**). With PGE2 pretreatment, hair matrix cells also decreased from 12 to 72 hours after IR, but the cell reduction was attenuated (**Fig.2D**). From 72 hours to 7 days, matrix cell numbers were progressively restored. These results showed that PGE2 pretreatment attenuated IR-induced apoptosis of hair matrix cells and increased cell proliferation, which contributed to the preservation of a higher matrix cell number.

### PGE2 pretreatment does not activate bulge stem cells for regeneration

Previous reports showed PGE2 promotes tissue regeneration following genotoxic injury through its effects on tissue stem cells [21, 23, 24]. Next, we determined whether PGE2 pretreatment promotes HFSC activation for regeneration. We performed continuous BrdU labeling from 12 hours after IR to 7 days after IR, when AHFR had been completed. Without PGE2 pretreatment, HFSCs in the bulge region were not labeled by BrdU (**Fig.3A**), indicating HFSCs did not actively proliferate during this period. Therefore, in the dystrophic catagen response induced by 8.5Gy of IR, HFSCs remained quiescent. In the PGE2 pretreatment group, no bulge HFSCs were labeled with BrdU either (**Fig.3A**). This finding indicated PGE2 pretreatment did not activate HFSCs to repair the damaged hair bulbs. We also examined whether 8.5Gy of IR induced HFSC apoptosis. We did not detect positive TUNEL staining in HFSCs in non-PGE2-treated and PGE2-treated HFs (**Fig.3B**). Previous study showed IR of up to 5.5Gy does not induce bulge HFSC apoptosis [11]. Here, at an even higher dose of 8.5Gy, HFSC apoptosis was not observed. The results showed HFSCs exhibit exceptional resistance to IR-induced apoptosis.

**Figure 3.**
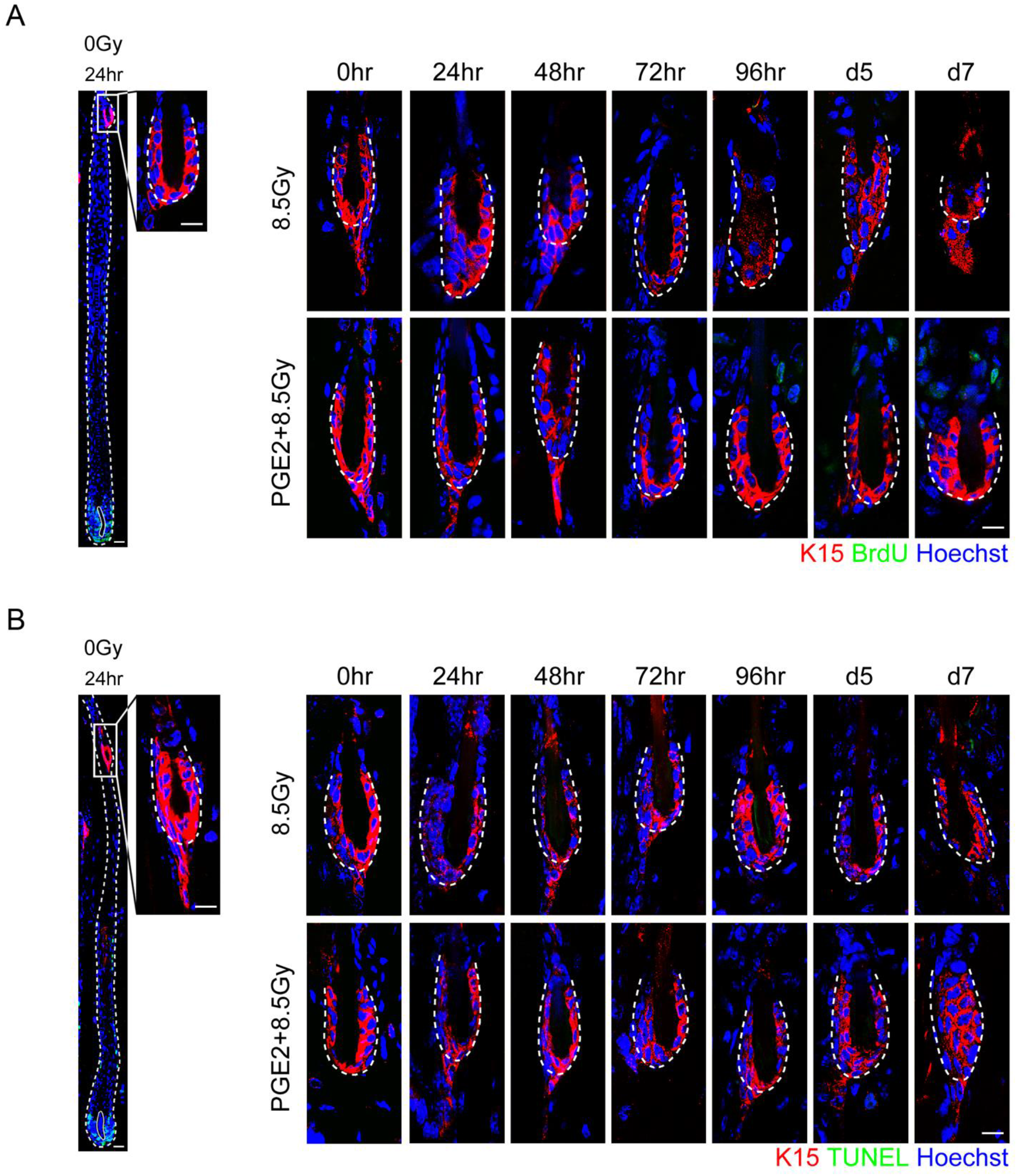
PGE2 pretreatment does not activate HFSCs and HFSCs are resistant to IR-induced apoptosis. Bulge HFSCs are stained by a K15 antibody. (**A**) Continuous BrdU labeling for analysis of HFSC proliferation. Left: low-power view of an unirradiated HF. Right: high-power view of the bulge region of irradiated HFs. No bulge HFSC is labeled with BrdU in both PGE2-treated and non-PGE2-treated groups. (**B**) TUNEL staining for analysis of HFSC apoptosis. Left: low-power view of an unirradiated HF. Right: high-power view of the bulge region of irradiated HFs. No TUNEL staining is detected in bulge HFSCs in both non-PGE2-treated and PGE2-treated HFs. Dashed line: basement membrane. Solid line: dermal papilla. Scale bar = 20 μ m in the low-power view, 10 μ m in the high-power view.

### PGE2 pretreatment transiently suppresses proliferation of matrix TACs

The delayed and reduced hair matrix cell apoptosis in the PGE2 pretreatment group suggests PGE2 protects matrix TACs from IR-induced apoptosis. Sensitivity to DNA damage from genotoxic IR varies across different phases of the cell cycle [32]. Cells are most sensitive to DNA damage-induced apoptosis in the G2-M phases and much less sensitive in the G1 phase. To test whether PGE2 pretreatment modulates the cell cycle of matrix TACs to protect them from IR-induced apoptosis, we analyzed the proliferation status of matrix TACs by BrdU pulse labeling before and after PGE2 pretreatment without IR. PGE2 treatment showed a time-dependent effect on matrix cell proliferation (**Fig.4A**). BrdU-labeled matrix cells became absent at 2 hours after PGE2 treatment and then gradually increased from 4 to 6 hours. The results suggested cell division of the highly proliferative matrix TACs transiently stopped or decreased from 2 to 4 hours after PGE2 treatment. Because we administered PGE2 2 hours before IR, IR injury to TACs fell within the time window when matrix TACs were proliferatively inactive. Therefore, PGE2 likely protects TACs from IR by reducing their radiosensitivity through changing their cell cycle phase, possibly to the G1 phase, when IR is delivered.

**Figure 4.**
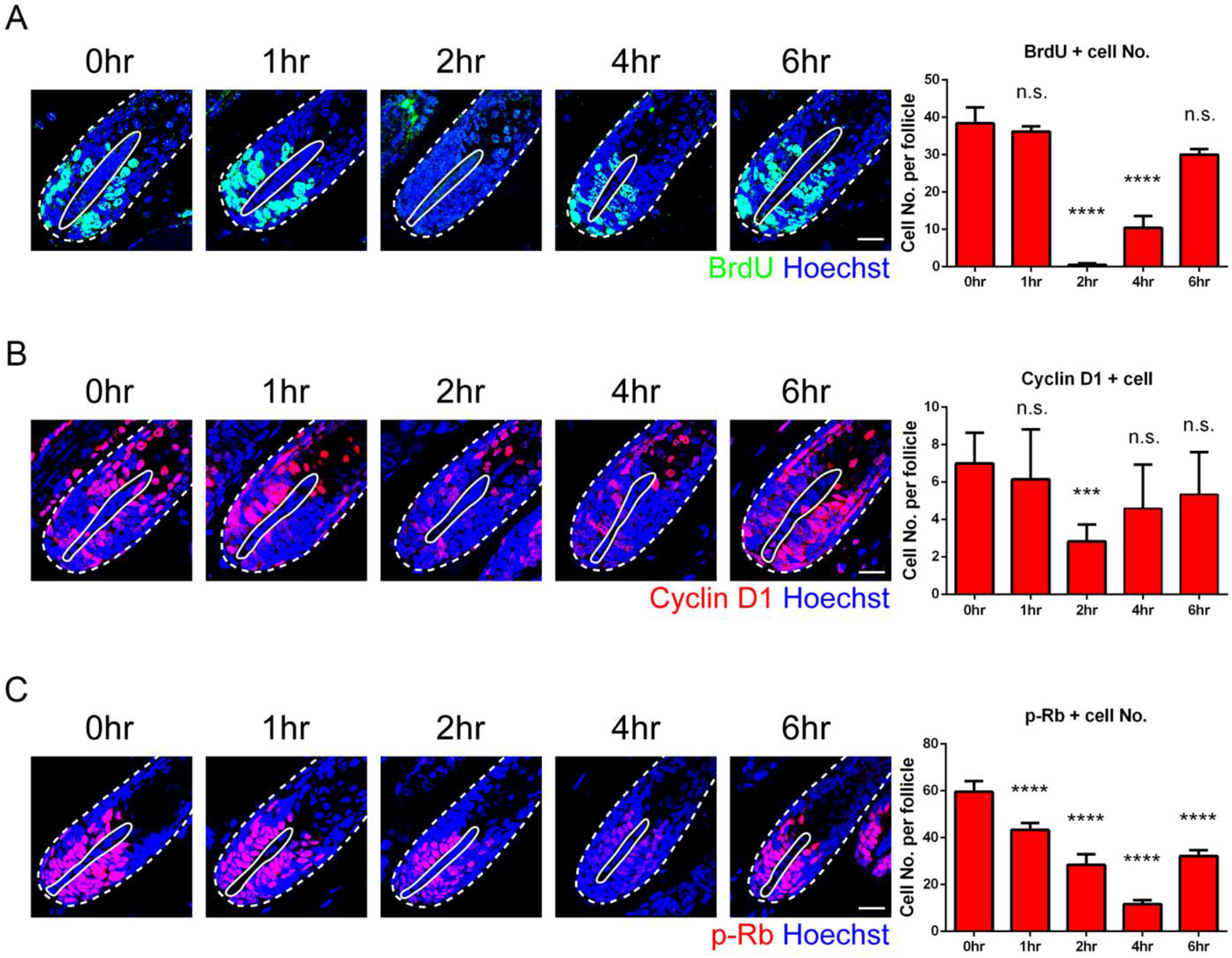
PGE2 transiently halts matrix TAC proliferation and suppresses cyclinD1 expression and Rb phosphorylation. (**A**) Pulse BrdU labeling before and after local PGE2 injection for analysis of matrix TAC proliferation. Numbers of BrdU positive cells in each hair bulb are quantified on the right side. (**B**) Immunofluorescent staining for Cyclin D1. Numbers of cyclin D1 positive cells in each hair bulb are quantified on the right side. (**C**) Immunofluorescent staining for pRb. Numbers of pRb positive cells in each hair bulb are quantified on the right side. Statistical significance was determined by Student’s *t* test. *** *p* < 0.001, **** *p* < 0.0001, n.s.: non-significant, compared with 0 hour. Error bars, mean + SEM. Dashed line: basement membrane. Solid line: dermal papilla. Scale bar = 20μm.

### PGE2 treatment decreases cyclin D1 and suppresses retinoblastoma tumor suppressor protein phosphorylation transiently

Cyclin D, retinoblastoma tumor suppressor protein (Rb), and E2F transcription factors control the G1 to S transition [33]. The unphosphorylated Rb actively inhibits cell cycle progression by binding the transactivation domain of the E2F transcription factor and recruiting chromatin-modifying enzymes to the promoters [34]. As the upstream cyclin D accumulates, it forms a complex with cyclin-dependent kinases 4/6 (CDK4/6)[35]. The cyclin D-CDK4/6 complex monophosphorylates Rb and, later, when Rb is hyperphosphorylated by CDK2, E2F is released and implements a transcriptional program that allows S-phase entry [33, 34]. Cyclin D and phosphorylated Rb (pRb) levels reflect the decision regarding G1 to S progression [33]. To determine whether PGE2 pretreatment arrests matrix TACs in G1 phase, we examined the abundance of cyclin D1 and pRb in HF TACs. Immunostaining of cyclin D1 showed a decreasing trend from 1 to 2 hours after PGE2 treatment, and then cyclin D1 showed an increase from 2 to 6 hours after PGE2 treatment (**Fig.4B**). pRb also showed a decreasing trend from 1 to 4 hours after PGE2 treatment and then increased (**Fig.4C**). The short suppressive effect of PGE2 on cyclin D1 and Rb phosphorylation suggests that PGE2 may halt TAC proliferation by transiently arresting them in the G1 phase. More matrix TACs are preserved when IR is delivered during the time window of G1 arrest when cells are less radiosensitive.

### PGE2 treatment leads to G1 arrest of HF epithelial cells

Next, we examined whether PGE2 induced G1 arrest in HF epithelial cells using the FUCCI reporter mice. FUCCI is a fluorescent reporter technology in which G1 and/or S/G2/M phases of the cell cycle can be determined using a dual-color scheme of red and green [29]. In the FUCCI system, the licensing factor Cdt1 level peaks in G1 and declines abruptly as S phase begins while its inhibitor geminin level is low during late mitosis and G1 phase, but high during S and G2 phases [36, 37]. As these FUCCI probes mark cells in G1 phase with red fluorescence and cells in S/G2/M with green, they can be used to identify living cells in either G1 or S/G2/M based on which fluorescent FUCCI probe they express. To monitor cell cycle changes over time, we cultured HF epithelial cells from FUCCI mice and used a time-lapse recording of fluorescence for 1 day after PGE2 treatment. We found green fluorescence (S/G2/M) cell numbers gradually decreased (**Fig.5A,5B**) in the presence of PGE2. In contrast, the red fluorescence (G1) cell numbers increased. This demonstrated PGE2 halts cell cycle progression of HF keratinocytes in G1 phase. This finding was consistent with the suppressive effect on cyclin D1 and Rb phosphorylation above.

**Figure 5.**
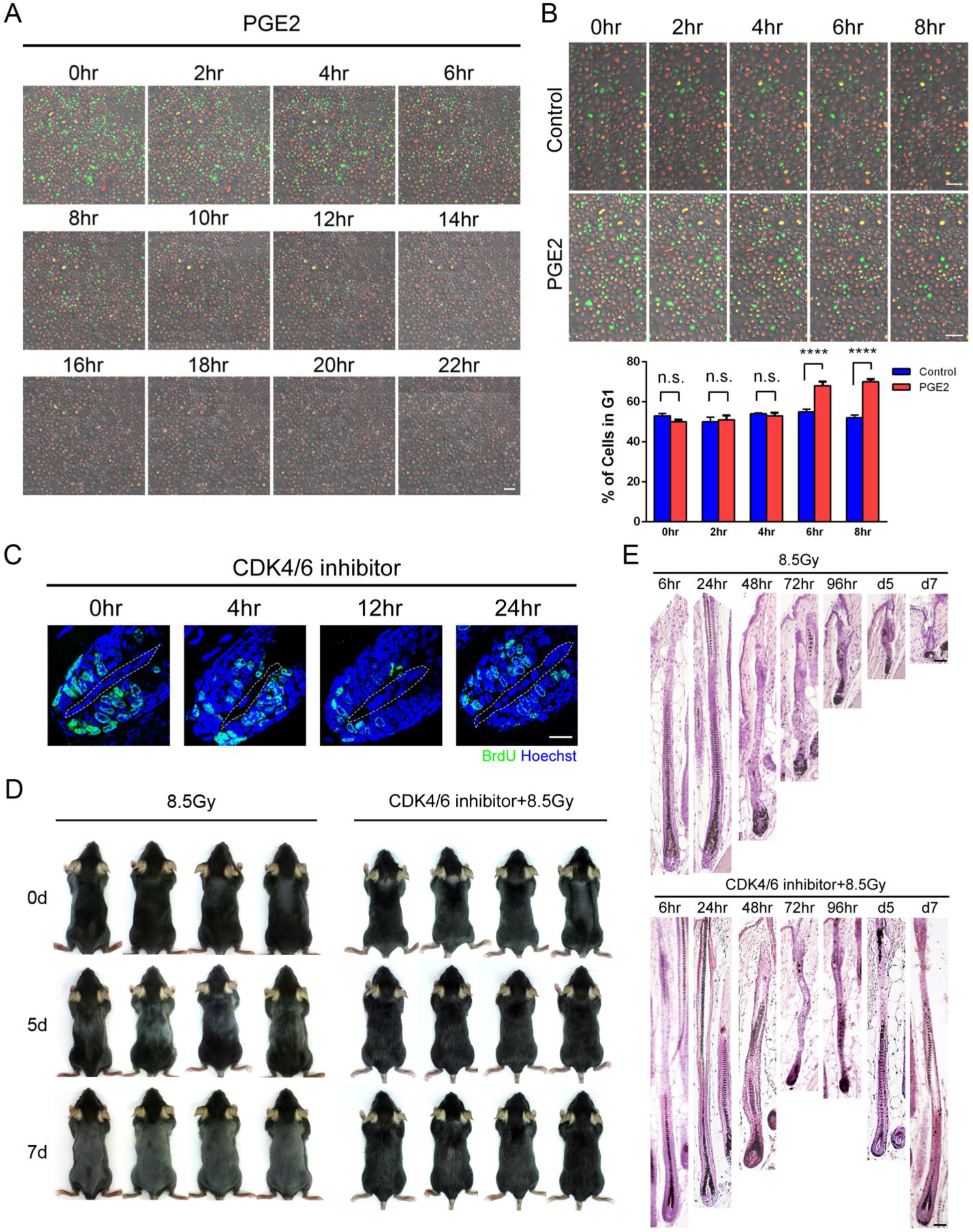
PGE2 treatment induces HF epithelial G1 arrest and pretreatment with a CDK4/6 inhibitor reduces IR-induced hair loss. (**A-B**) Effect of PGE2 on cell cycle progression of cultured HF epithelial cells. Time-lapse recording of the fluorescence of HF epithelial cells from FUCCI reporter mice. In the presence of PGE2, cells with green fluorescence (S/G2/M) gradually decreased in the presence of PGE2 while cells with red fluorescence (G1) increased. Compared with control, the PGE2-treated group has higher percentage of cells in G1 phase (red fluorescence) at 6 and 8 hours. (**C**) Pulse BrdU labeling before and after CDK4/6 inhibitor, Palbociclib Isethionate (PD0332991), treatment. Cell proliferation of matrix TACs was highly suppressed at 12 hours and increased back at 24 hours post the treatment of PD0332991. (**D**) Gross pictures of mice with or without PD0332991 pretreatment. Pretreatment with PD0332991 12 hours before IR reduces hair loss. (**E**) Histology of HFs with PD 0332991 pretreatment. Dystrophy of HFs is reduced and HF repair is accelerated in the PD0332991-treated mice. Statistical significance was determined by Student’s t test. \*\*\*\**p* < 0.0001, n.s.: non-significant. Error bars, mean + SEM. Dashed line, DP. Scale bar = 100 μ m in A, B; 20 μ m in C, 100 μ m in E.

### Induction of TAC G1 arrest by the CDK4/6 inhibitor prevents IR-induced hair loss

The results above suggest PGE2 pretreatment reduces the radiosensitivity of HF TACs by arresting their cell cycle in the G1 phase. To mimic this, we tested the effect of palbociclib isethionate (PD0332991), a CDK4/6 inhibitor, on matrix TACs. We found that the proliferation of matrix TACs was highly suppressed at 12 hours and increased at 24 hours after PD0332991 treatment (**Fig.5C**). This finding suggested more matrix TACs were in G1 phase at 12 hours after PD0332991 treatment. Therefore, we pretreated mice with PD0332991 12 hours before exposure to 8.5Gy of IR. PD0332991 pretreatment yielded a protective effect on RIA loss similar to that of PGE2 (Fig.5D). In contrast to premature termination of anagen in mice without PD0332991 pretreatment, HF dystrophy was reduced, and HF repair was accelerated with PD0332991 pretreatment (Fig.5E). Restoration of the anagen morphology on day 7 after IR demonstrated a successful reparative response. Therefore, the transient induction of G1 arrest by the CDK4/6 inhibitor also prevented RIA by protecting matrix TACs and enhancing HF repair.

## Discussion

In this study, we showed that local PGE2 pretreatment can prevent RIA. This protective effect is mediated not by activating HFSCs for regeneration, but by attenuating HF dystrophy through reducing the radiosensitivity of HF TACs. Local administration of PGE2 transiently arrests rapidly dividing HF TACs in the G1 phase at about 2–4 hours after PGE2 treatment, during which radiosensitivity is diminished. TAC apoptosis was reduced when IR was administered during this time window. PGE2 pretreatment reversibly downregulates cyclin D1 and reduces Rb phosphorylation in HF TACs, resulting in transient G1 arrest. Preservation of more TACs reduces HF dystrophy, prevents premature termination of the anagen phase by bypassing entry into the catagen/telogen phase, and accelerates HF repair for structural restoration.

In response to genotoxic RT and chemotherapy, anagen HFs undergo either a dystrophic anagen response or a dystrophic catagen response according to the severity of injury (**Fig.6**) [11, 14, 17, 38, 39]. Anagen HFs can elicit an AHFR scheme in response to injuries to restore the lost structures to resume anagen growth [11, 17]. In the dystrophic anagen response, usually after a relatively limited injury, the anagen HF structure is successfully restored. In contrast, when HFs are damaged more extensively, AHFR fails to regenerate the lost anagen structure, and HFs prematurely enter catagen and then telogen, i.e., the dystrophic catagen response. Hair regrowth is not observed until a long delay, after which HFs reenter a new anagen fueled by the activation of HFSCs. Here, we demonstrated that PGE2 pretreatment changes the decision of entering dystrophic catagen to dystrophic anagen (**Fig.6**). Because this conversion bypasses the long downtime needed to rest the hair cycle through telogen for HFSC activation, hair loss is much attenuated.

**Figure 6.**
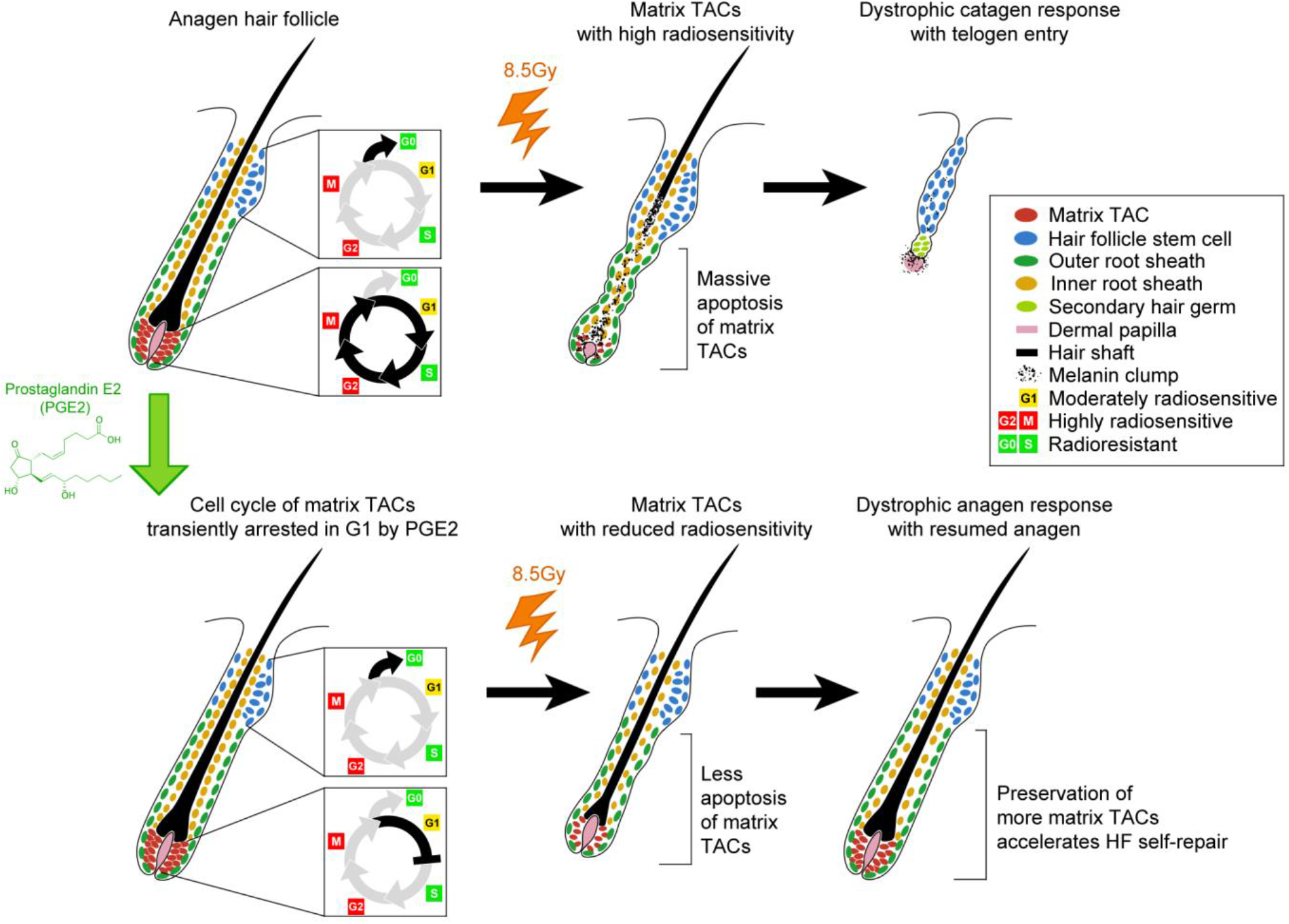
Prostaglandin E2 prevents radiotherapy-induced alopecia by attenuating TAC apoptosis through promoting G1 arrest. Depiction of the protective effect of PGE2 pretreatment on IR-induced HF damage

After radio- or chemotherapeutic injury, we showed that the lost TACs are mainly replenished during AHFR by the basal lower proximal cup and outer root sheath cells, which have a plastic stem cell-like property, and their progeny become new TACs [11, 17, 39]. Because PGE2 pretreatment preserves more TACs after IR, it reduces the number of TACs that need to be replenished by local progenitors, thus accelerating the regenerative process. Since preserving more TACs can alter the regenerative dynamics toward a successful AHFR, the decision to undergo either a dystrophic anagen response or a dystrophic catagen response by the HF is modulatable. This highlights an important strategy to prevent RIA by tilting the balance toward successful AHFR to avoid premature catagen entry by maintaining a critical number of TACs. As the cellular dynamics for AHFR after chemotherapy is similar to that after RT [11, 17, 39], it is highly possible that this strategy can be applied to the prevention of chemotherapy-induced alopecia.

Strategies of modulating cell cycle for normal tissue protection from genotoxicity are emerging [40–42]. Using a CDK4/6 inhibitor to induce G1 arrest attenuates IR injury to the intestine by reducing apoptosis of the proliferating Lgr5^+^ stem cells [41, 43, 44]. Palbociclib, a CDK4/6 inhibitor, protects human HFs from chemotherapy-induced damage ex vivo by arresting HFs in G1 phase [45]. Here we showed that PGE2 arrests HF TACs in G1 phase in vivo in a manner similar to that of the CDK4/6 inhibitor. Because traditional radioprotectors can potential provide indiscriminate protection to both normal tissues and cancer cells [46], it is crucial to limit cell cycle modulation to the desired normal tissues, either by targeting tissue-specific pathways or through tissue-specific delivery of a cell cycle modulator. From this perspective, the transient effect of PGE2 to function locally for efficient HF protection provide a unique advantage for clinical application. PGE2 can be administered 2–4 hours before RT by multiple intracutaneous injections or arrays of fine needles across the target skin region to limit its systemic effect.

The mechanism by which G1 arrest attenuates radiosensitivity remains unclear. Our results of γ-H2AX showed PGE2 pretreatment significantly reduced foci of DNA damage in the hair matrix after IR (**Fig.2E**). Since more TACs are in G1 phase after PGE2 pretreatment, there is less DNA content per cell due to the halt in active DNA replication. With less DNA exposed to IR, it is reasonable that the average foci of DNA breaks per TAC induced by a similar IR dose would be reduced. Another possibility is that the efficiency of DNA repair can vary among different phases of the cell cycle [47]. If the efficiency of DNA repair is enhanced in G1 phase, this can also reduce the radiosensitivity of TACs. It is also notable that apoptosis is not only reduced but also delayed after PGE2 pretreatment. Compared with prominent TAC apoptosis initiated at 6 hours after IR in the PGE2 pretreatment group, TAC apoptosis was not observed until 12 hours after IR (**Fig.2B**). Therefore, PGE2 pretreatment might not only change the extent of DNA breaks and the subsequent DNA repair dynamics but also alter the signaling network for the induction of apoptosis. The delay in apoptotic induction allows cells to repair more DNA damage before making the decision to initiate apoptosis. Therefore, the protective effects of PGE2 on TACs may result from a combinatorial effect.

## Conclusions

In this study, we show PGE2 pretreatment attenuates RIA by reducing apoptosis of HF TACs. Locally administered PGE2 transiently arrests HF TACs in G1 phase and thus reduces their radiosensitivity. When IR is delivered in this time window of higher radioresistance, more TACs are protected from IR-induced apoptosis. Preservation of more TACs accelerates AHFR, bypasses the hair cycle resting through catagen/telogen and thus reduces hair loss. Because PGE2 is currently used clinically for other treatment, it can be formulated and repurposed for local skin administration to prevent RIA. It can also potentially prevent hair loss from genotoxic chemotherapy.

## Acknowledgments

This work was supported by Taiwan National Science and Technology Council (MOST106-2314-B-002-133-MY3, MOST109-2314-B-002-072-MY3 to SFL), National Taiwan University Hospital (108-N4174, 109-N4525, 109-O23 to SFL; 111IF0006 to SJL), Taiwan National Health Research Institutes (NHRI-EX111-1112EI to SJL) and National Taiwan University (111L892401-04 to SJL). The authors thank the 2nd and 8th Core Lab, Department of Medical Research, National Taiwan University Hospital, for technical support and Dr. Hironobu Fujiwara for the help in obtaining FUCCI mice. We also thank the members in SJL laboratory for their discussion

