## Supplementary material for "Prostaglandin E2 prevents radiotherapy-induced alopecia by attenuating transit amplifying cell apoptosis through promoting G1 arrest": supplement.pdf

### Supplementary methods

#### Image acquisition and quantification

All fluorescent images were acquired using a confocal microscope (SP8; Leica, Wetzlar, Germany). We acquired 1,024 × 1,024 pixels sequential scans with a 63X oil-immersion objective lens (1.4 NA) to quantify BrdU-positive and TUNEL positive cells. The hair matrix cells were counted as HF epithelial cells below the top of the dermal papilla.

#### Cell culture and time-lapse images

Primary HF epithelial cells were isolated from the vibrissae of FUCCI mice and cultured in a serum-free defined keratinocyte serum-free medium (KSFM). For cell cycle analysis, vibrissal HFs were isolated from FUCCI mice, and the lower hair bulbs containing dermal papillae were removed using a scalpel. The upper part of the vibrissal HFs was cultured in an imaging dish (ibid, Cat. 81156) with serum-free defined KSFM. After 2 weeks of culture, PGE2 was applied to the cultured keratinocytes, and time-lapse images were acquired to track cell cycle dynamics. Time-lapse images were acquired using a Leica SP8 confocal microscope at an interval of 5 minutes. Green and red fluorescencex were excited at 488 nm and 540 nm, respectively, by laser. Emission was captured through band-pass settings of 505–550 nm and 590–620 nm.

#### Quantification of r-H2AX foci

DNA double-strand breaks were determined using γ-H2AX foci. The 3-dimensional fluorescent images were reconstituted by approximately 10–15 serial two-dimensional Z-stacks of confocal images [32]. Foci within Hoechst-stained nuclei were scored based on the value of pixels after the focus threshold was manually set as we previously described[11]. We compared the total number of positive pixels within the nucleus in selected cells to estimate the number of foci in the entire nuclear region as we previously described [11].

#### Statistical analysis

Data are presented as the mean ± standard error of the mean. The unpaired Student *t*-test was used to compare datasets between the two groups of mice. To compare three or more groups, we performed one-way analysis of variance followed by Bonferroni multiple comparisons. Statistical comparisons were performed using the Prism software (GraphPad Software, Inc., San Diego, CA). Statistical significance was set to $p < 0.05$.

**Supplementary Table S1. Antibodies used**

| **Antibody** | **Source** | **Catalog number** | **Dilution** |
|---|---|---|---|
| BrdU | Abcam | ab6326 | 1/200 |
| γ-H2AX | Millipore | #MABE205 | 1/100 |
| Cleaved caspase 3 | Cell Signaling | #9664 | 1/200 |
| Cyclin D1 | Abcam | ab16663 | 1/200 |
| Phosphorylated Rb (pRb) | Cell Signaling | #8516 | 1/100 |
| Cytokeratin 15 | Thermo | MA1-90929 | 1/100 |

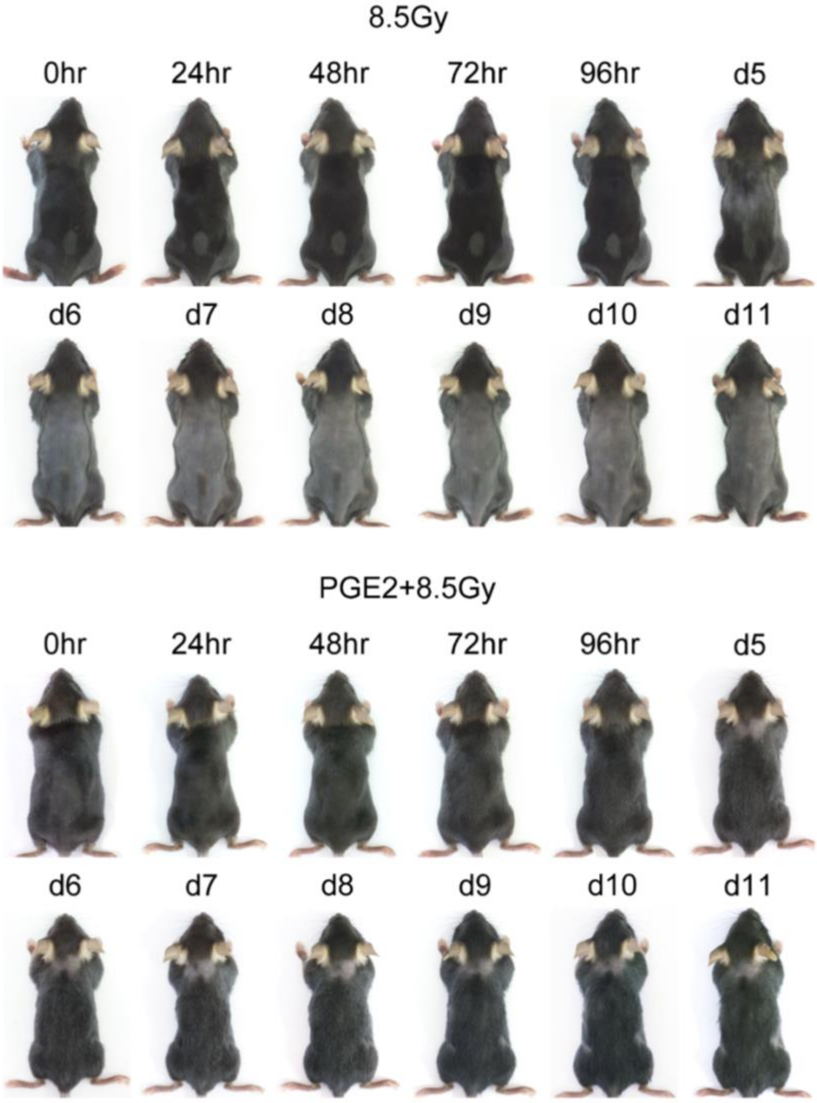


**Supplementary Figure 1. Gross pictures of mice with and without local PGE2 pretreatment before IR.** IR of 8.5Gy induced prominent hair loss in mice without PGE2 from day 5 post IR. With local PGE2 pretreatment 2 hours before IR, hair loss was pronouncedly reduced.
